# Human cortical dynamics during full-body heading changes

**DOI:** 10.1101/417972

**Authors:** Klaus Gramann, Friederike U. Hohlefeld, Lukas Gehrke, Marius Klug

## Abstract

The retrosplenial complex (RSC) plays a crucial role in spatial orientation by computing heading direction and translating between distinct spatial reference frames based on multi-sensory information. While invasive studies allow investigating heading computation in moving animals, established non-invasive analyses of human brain dynamics are restricted to stationary setups. To investigate the role of the RSC in heading computation of actively moving humans, we used a Mobile Brain/Body Imaging approach synchronizing electroencephalography with motion capture and virtual reality. Data from physically rotating participants were contrasted with rotations based only on visual flow. During physical rotation, varying rotation velocities were accompanied by pronounced wide frequency band synchronization in RSC, the parietal and occipital cortices. In contrast, the visual flow rotation condition was associated with pronounced alpha band desynchronization, replicating previous findings in desktop navigation studies, and notably absent during physical rotation. These results suggest an involvement of the human RSC in heading computation based on visual, vestibular, and proprioceptive input and implicate revisiting traditional findings of alpha desynchronization in areas of the navigation network during spatial orientation in movement-restricted participants.

## Introduction

Heading computation is fundamental for spatial orientation in humans and other species. The registration of moment-to-moment changes in orientation with respect to an allocentric (view-point independent) reference direction provides information about an animal’s current heading relative to the environment. This is accomplished by the integration of vestibular, proprioceptive, and visual signals providing information about linear and angular velocity signals of the head, the relative position of the head with respect to the trunk, and information about stable aspects of the environment, respectively (Angelaki & Cullen, 2008). Single cell recordings in freely behaving animals identified several brain structures involved in heading computation, including the retrosplenial cortex (Chen et al., 1994; Cho &Sharp, 2001). The RSC receives input from the visual system and from head direction cells in the thalamic nuclei (Vann et al., 2009). It also hosts subpopulations of heading-sensitive cells that are sentient to local features of the environment, while other cells exhibit mixed activity patterns related to both local and global heading computation (Jacob et al., 2017). These findings suggest that neural activity in the RSC subserves the integration of egocentrically (view-point dependent) coded landmark cues based on sensory fusion (vision and proprioception; Fischer et al., 2020; Mitchell et al., 2018) with allocentric heading information originating from the Papez circuit (Taube, 1998). This allows the compensation of rotational offsets between egocentric and allocentric spatial representations, routed from the parietal and medial temporal cortices, providing the necessary information for translating between both frames of reference in the RSC (Byrne et al., 2007; Ekstrom et al., 2017).

The central role of the RSC for spatial orientation in general and for heading computation specifically is supported by human imaging studies (Epstein, 2008; Maguire, 2001; Mitchell et al., 2018). Due to the restricted anatomical differentiation of the retrosplenial (BA 29 and 30) and the adjacent posterior cingulate cortex (BA 23 and 31), the abbreviation RSC is used here to refer to the retrosplenial complex (Chrastil, 2018; Epstein, 2008). Haemodynamic changes in the RSC were shown to be associated with landmark learning (Auger et al., 2012; Spiers &Maguire, 2006), with both global and local heading estimation (Marchette et al., 2014), and with translating view-point independent location representations into behaviourally relevant egocentric representations (Berens et al., 2021). However, while functional magnetic resonance imaging (fMRI) studies provide valuable insights into the function of the RSC regarding spatial cognition, they do not allow movements of the participant in the scanner (Gramann et al., 2011). This is due to the fact that fMRI studies use sensors that are too heavy to follow movements of the signal-generating source (Gramann et al., 2014). Electroencephalography (EEG) studies, though utilizing lighter sensors, are considerably affected by movement-related artefacts and thus traditionally rely on stationary setups as well. Crucially, heading computation depends on input from the vestibular organ (Angelaki &Cullen, 2008) indicating movement of the head and body that can be related to, among other features, the location and orientation of external information like landmarks encoded through other senses (Jeffery et al., 2016). Therefore, established imaging studies do not allow a recording of the very signal that is essential for heading computation, fostering cognitive processes in established brain imaging studies that might not resemble the computation and use of directional heading in more natural environments (Gramann, 2013).

In the present study, we overcame restrictions of traditional imaging studies by investigating neural dynamics in the human RSC during heading computation in actively rotating humans. To this end we used a Mobile Brain/Body Imaging approach (Gramann et al., 2011; Makeig et al., 2009) synchronizing high-density EEG to motion capture and head-mounted virtual reality (VR). Data-driven analyses, based on spatial filtering and subsequent source reconstruction, were used in order to investigate neural dynamics and their neuroanatomical origins accompanying heading computation during physical rotations in movement-unrestricted participants. The analyses focused on the RSC as the cortical structure which was suggested to be involved in heading computation, and additionally the occipital and parietal cortex reflecting visual processing of heading changes as well as multisensory fusion of vision and proprioception, respectively. The results demonstrate significant spectral modulations in a wide frequency range in the RSC as well as the occipital and parietal areas during active physical rotations compared to a stationary setup that provided only visual flow.

## Results

We analysed data from 19 participants performing a spatial orientation task in two rotation conditions (see Figure 1A–C; see supplement for videos of the setup), involving i) physical rotations of the whole body (“physR”), and ii) standing in front of a desktop monitor controlling the visual flow by manually operating a joystick (“joyR”). The latter condition replicated traditional stationary setups investigating neural dynamics underlying spatial orienting based only on visual flow. The participants rotated on the spot in a sparse virtual environment that provided an initial local landmark (pole) but no other stable features. With a button press, the landmark was replaced by a sphere, which moved either to the left or to the right around the participant at a constant distance. In this outward rotation phase, participants had to follow the movement of the sphere by rotating on the spot using the respective control (physR or joyR). The sphere followed one of two possible cosine velocity profiles along its path and stopped at varying eccentricities with respect to the participants’ initial facing direction. Upon stopping, the sphere changed its colour, and the participants were tasked with rotating back and indicating their initial heading. EEG data was decomposed into independent components (ICs) using adaptive mixture independent component analysis (AMICA; Palmer et al., 2008). The approximate locations of the resultant ICs were reconstructed using equivalent dipole models, and the ICs were clustered using a repetitive k-means algorithm optimized to the RSC as a region of interest (see Methods for details).

**Figure 1:**
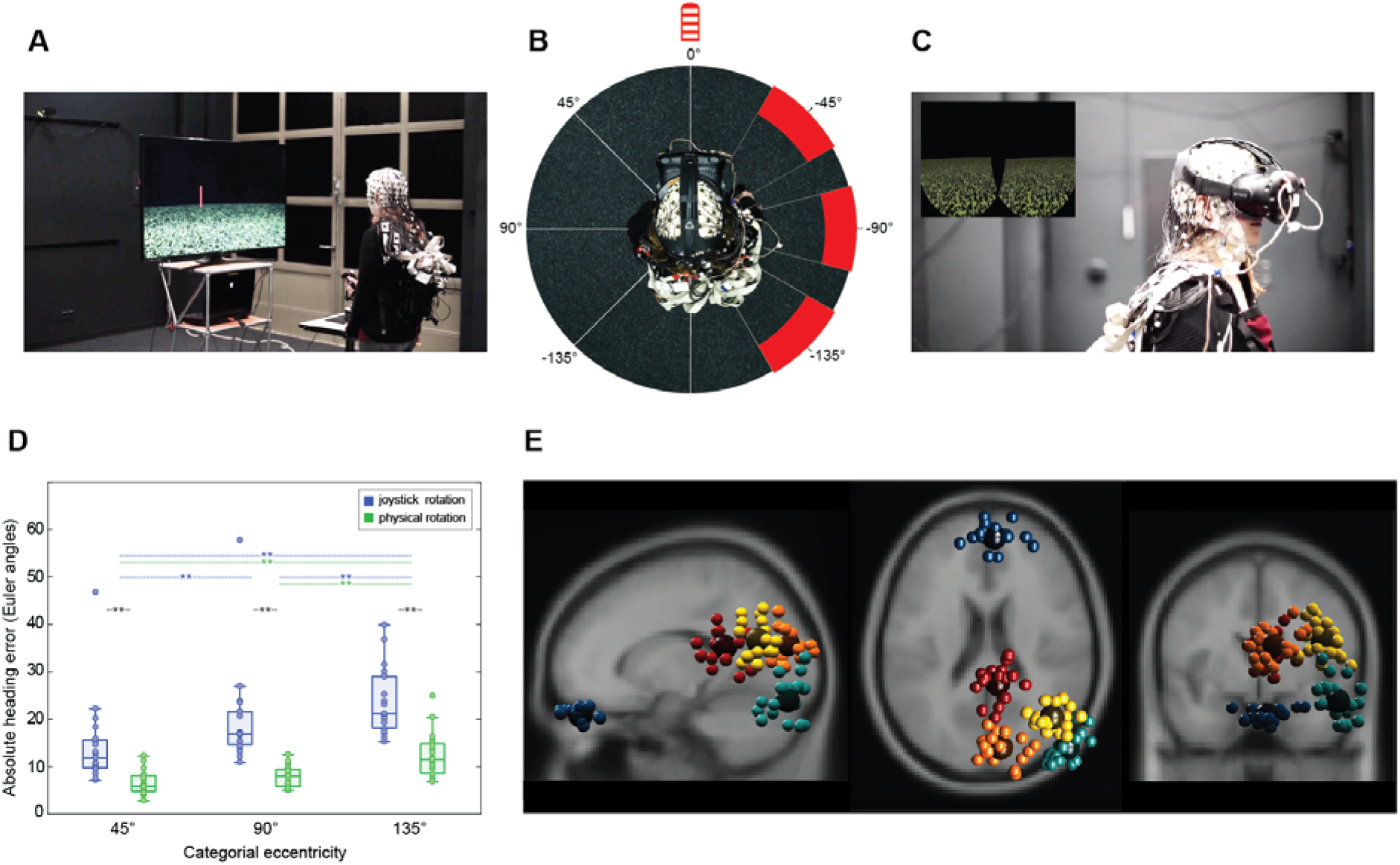
Experimental setup, heading error and representative IC-cluster. **A**) Setup of the stationary condition with joystick rotation (joyR; visual flow only), displaying a sparse virtual environment with a local landmark providing the initial heading direction (pole). The joystick was placed on a table in front of the standing participant. **B**) Top-down view of a participant in the physical rotation (physR) condition with MoBI setup, displaying the rotation eccentricities (categorial eccentricities varying +/-15° around 45°, 90°, and 135°, respectively). **C**) MoBI setup with a participant wearing high-density EEG synchronized to motion capture (red LEDs on VR goggle) and a head-mounted VR. The inset displays the binocular view of the virtual environment. **D**) Absolute heading error (orientation yaw; Euler angles) after completing the back rotation, displayed for both rotation conditions as a function of eccentricity, averaged across rotation directions. The boxplot comprises all participants (median; whiskers extending to 1.5 times the interquartile range). Bonferroni-significant p-values of post hoc testing are shown (Wilcoxon signed-rank test). ** indicates p < 0.01. **E**) Representative clusters of independent components (ICs) with single ICs displayed as small spheres and cluster centroid displayed as larger spheres. ICs are projected onto a standard brain (MNI) with sagittal, horizontal and coronal views from left to right. Cluster centroid in Talairach space for a cluster representing eye movement activity (blue; x=4, y=46, z=−28; no BA); a cluster representing right neck muscle activity (light blue; x=54; y=−85, z=−10; no BA); a cluster representing activity originating in or near the restrosplenial complex (RSC) (dark red; x=8, y=−42, z=18; BA30); a cluster representing activity originating in or near the right inferior parietal cortex (yellow; x=44, y=−63, z=23; BA39); a cluster representing activity originating in or near the occipital cortex (orange; x=9, y=−81, z=20; BA18).

### Heading estimation is more accurate for physical rotation

Replicating previous results, the performance data showed that physR resulted in higher accuracy for heading reproduction than joyR (Jürgens et al., 1999; Klatzky et al., 1998). A repeated measures analysis of variance (rANOVA) for the mean absolute heading error, averaged over categories of pseudo-continuous eccentricities of 3° steps centred around 45° (30–60°), 90° (75–105°), and 135° (120–150°), revealed a significant main effect of “rotation condition” (F_1,18_ = 33.78; p < 0.001; partial η^2^ = 0.65) and of “eccentricity” (F_1.28,23.11_ = 26.59; p < 0.001; partial η ^2^ = 0.6), as well as a significant interaction between both factors (F_1.78,31.97_ = 5.75; p = 0.009; partial η ^2^ = 0.24). Post hoc analysis of the interaction effect using the Wilcoxon signed-rank test revealed that the absolute heading error was significantly smaller for the physR than for the joyR condition in all three eccentricity categories (45°: 6.76 ± 2.77 vs. 14.4 ± 8.9; 90°: 7.79 ± 2.21 vs. 19.73 ± 10.16; 135°: 12.38 ± 4.68 vs. 23.83 ± 7.1; p < 0.001 for all comparisons). The post hoc tests also revealed an increase of absolute heading errors with increasing eccentricity in the joyR condition (p < 0.01 for all comparisons), but significant differences in the physR condition only for the most eccentric positions of 135° as compared to both 45° and 90°.

### Head rotation velocity differentiates oscillatory neural activity

To test whether variable angular movement information from the vestibular and visual systems is associated with changes in the oscillatory amplitudes (Hilbert-transform; Clochon et al., 1996) of theta, alpha, and beta frequency bands, we applied single trial movement velocity binning (Bassett &Taube, 2001; Linkenkaer-Hansen et al., 2004) and Mahalanobis distance-based Representational Similarity Analysis (RSA; Kriegeskorte &Kievit, 2013; Nili et al., 2014) to both the physR and joyR rotation conditions. Briefly, movement velocity (yaw orientation) during the outward rotation following the visual stimulus was extracted from movement onset to offset for all trials, and subsequently a velocity binning procedure was applied (cf. Supplements for details), as previously established for single cell recordings of heading-sensitive cells in rodents (Bassett &Taube, 2001) and in the context of analysing EEG oscillations (Linkenkaer-Hansen et al., 2004). The binning analysis resulted in 10-percentile velocity bins (ranging from slowest to largest movement velocity), and the baseline-corrected oscillatory amplitudes in each IC were averaged per velocity bin separately for each frequency band (9 bands; non-overlapping 2.5 Hz steps from 4–30.5 Hz) and across all trials, thus obtaining a single value per participant IC and velocity bin. Subsequently, within the RSA framework (Nili et al., 2014) the Mahalanobis distance (Mahalanobis, 1936) was calculated for all combinations of frequency band × velocity bin × rotation condition, thus obtaining a 180 × 180 Representational Dissimilarity Matrix (RDM) for each participant’s IC in a given cluster. Significance across participants in selected clusters was obtained by permutation-based statistics (p=0.05; cf. Supplements for details). Grand-averages and statistical results are presented in Figure 2.

**Figure 2.**
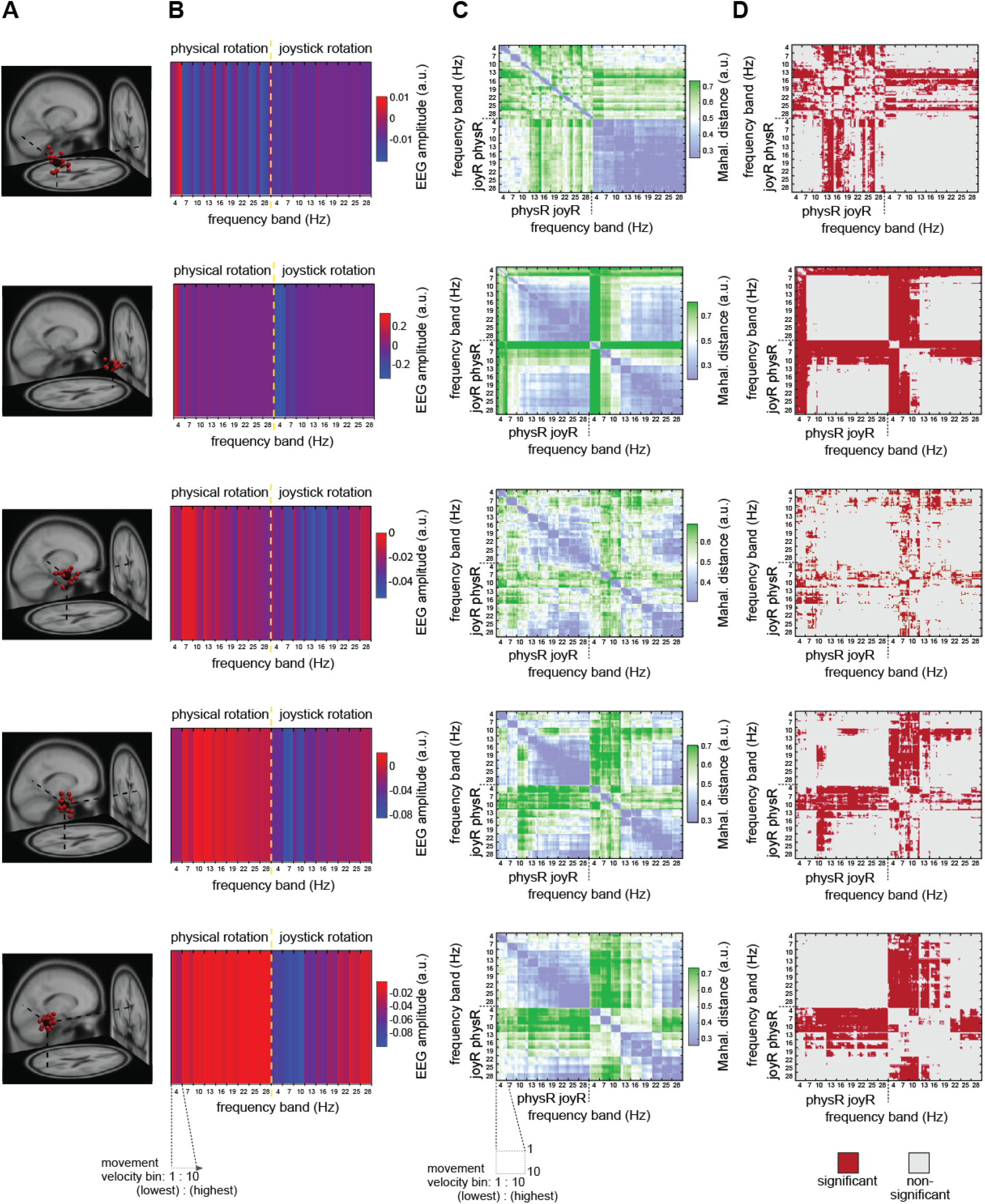
Representational Similarity Analysis (RSA) of movement velocity-associated modulation of oscillatory activity in the right neck cluster. **A)** 3D-projection of IC-clusters onto a standard brain (described in detail in figure 1). **B)** Grand-average IC amplitudes across all ICs in the cluster sorted according to velocity bins from lowest velocity to highest velocity; color bar is scaled to min. and max. Displayed are the start categories of each frequency band (9 bands; non-overlapping 2.5 Hz steps), and each frequency band contains, in ascending order, 10 movement velocity bins (percentiles; 10-100 % referring to slowest and largest velocities, respectively). **C) G**rand-average normalized Mahalanobis distance (Representational Dissimilarity Matrix, RDM); color bar is symmetrically scaled to 85% of the max. value; the normalized Mahalanobis distance scales from 0 (no distance) to 1 (max. distance), values of ∼0.5 are obtained on randomly shuffled data. **D)** RDM statistical significance (tested vs. noise level, permutation testing with n=10000 permutations, p=0.05) for the outward rotation (movement onset to offset). FDR – false discovery rate; IC – independent component; RDM – representational dissimilarity matrix; RSA – representational similarity analysis.

For the cluster representing activity of right-lateralized neck muscles, increasing grand average IC amplitudes were visible with increasing rotation velocities across a wide range of frequencies, most pronounced for the theta and beta bands, visible in the grading of grand average amplitudes within narrow-filtered frequency bands dependent on the velocity (Figure 2, first row). For the IC cluster representing eye movements, in contrast, decreased grand average amplitudes with increasing movement velocity, restricted to the theta frequency range, were observed (Figure 2, second row). Significant differences in movement velocity-associated modulation of oscillatory activity was only observed for the physR condition with increasing Mahalanobis distance with increasing movement velocity for the neck muscle cluster (beta range, 13–27.5 Hz). In contrast, for the eye cluster, both movement conditions revealed velocity-associated modulations (theta range, 4–6.5 Hz), as indicated by grading (striping) of the distance per frequency band from light to intense green coloring (Figure 2 statistical tests in Figures D). In the brain clusters (RSC, right parietal, occipital) both joyR and physR conditions showed velocity dependent variations in Hilbert-transformed amplitudes throughout the entire frequency range with decreasing amplitudes associated with increasing velocities and with pronounced differences between the movement conditions. These differences were also visible in the significant velocity-associated grading of the Mahalanobis distance (larger velocity ≈ larger distance). This effect, however, varied according to the topography and frequency bands: in joyR, increasing distance was observed in all three clusters in broad frequency ranges of approx. 4–21.5 Hz (theta, alpha bands: RSC, parietal, occipital; beta: RSC, occipital), whereas for physR the effect was pronounced only in the alpha (RSC, 7–9.5 Hz) and alpha/beta frequency ranges (parietal: 10–15.5 Hz) without significant velocity-associated distance modulations in the occipital cortex.

### Event-related spectral dynamics differ between physical and visual flow only rotation

To further investigate modulations of oscillatory neural activity during heading computation based on active body rotations as compared to visual flow only we investigated spectral dynamics during outward rotations. For this purpose, in a first step single-trial spectrograms were computed for all trials. In order to account for different trial durations associated with variable eccentricities, the spectrograms were linearly time-warped to the onset of the visual stimulus and to the movement onset as well as the movement offset at the trial end (participant’s head or joystick movement), resulting in time-warped event-related spectral perturbations (ERSP; see Figure 3). The spectral baseline was defined as the 200ms period before stimulus onset, excluding movement-contaminated trials. Here, brain activity originating from the RSC, as well as the parietal cortex and the occipital cortex, respectively are further discussed. In addition, the two non-brain clusters representing activity stemming from right neck muscle activity and eye movements are displayed to allow for comparison with brain dynamics.

**Figure 3:**
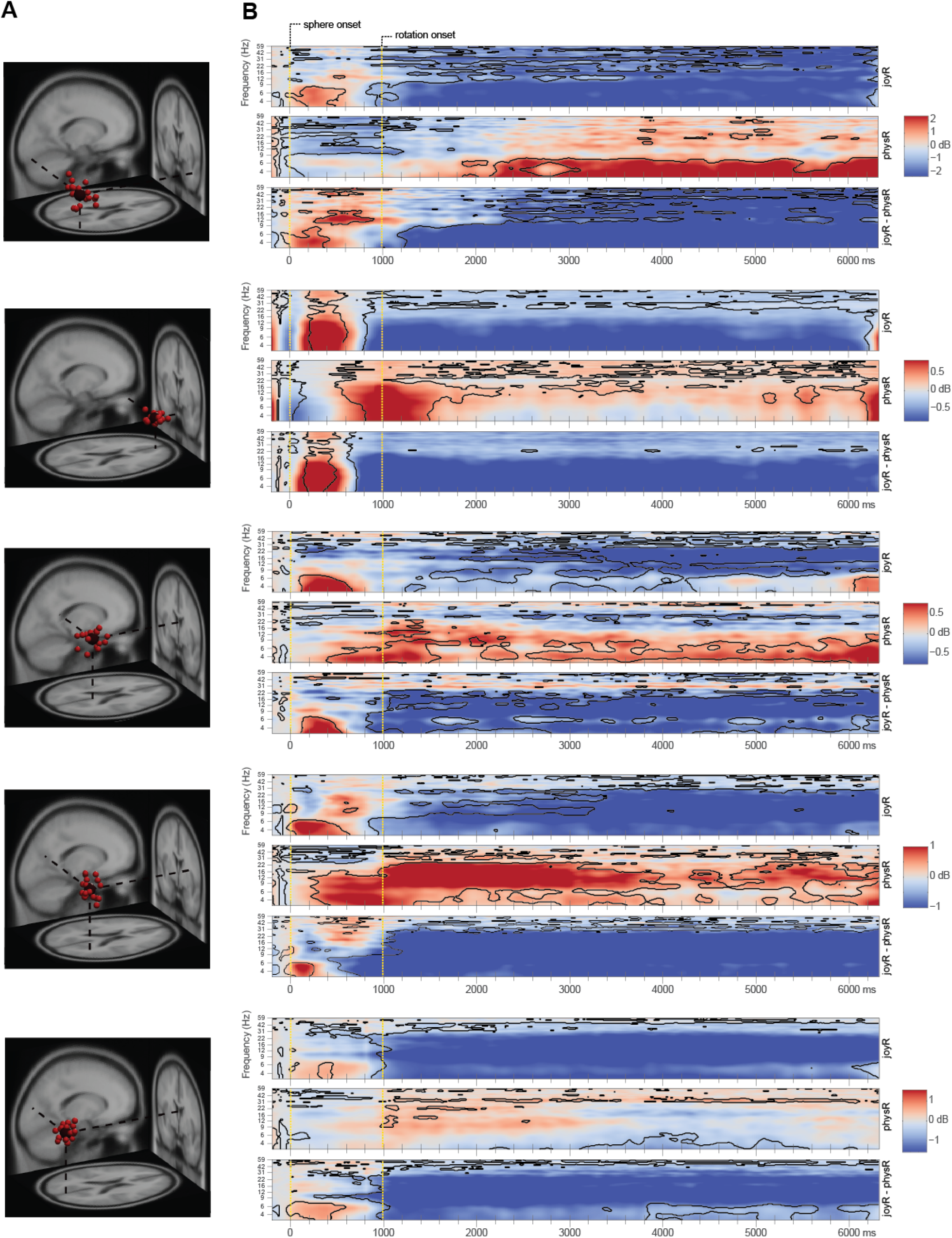
Event-related spectral perturbations (ERSPs) in representative IC-clusters. **A)** Clusters of ICs projected onto a standard brain space (MNI) with each small sphere representing individual ICs and the bigger sphere representing the cluster centroid (described in detail in figure 1). **B)** Time-warped event-related spectral perturbations (ERSPs) in different clusters. Epochs were time-warped with respect to the sphere stimulus (time point zero) and to the mean rotation onset (head or joystick movement; second dotted vertical line) as well as the movement offset (end of trial). Upper and middle rows of each time-warped ERSP: FDR-significant (0.01) differences to the baseline (−200ms to stimulus onset) are indicated by the traces around the respective time-frequency bins. **Upper row)** ERSP for the joystick rotation condition (joyR). **Middle row)** ERSP for the physical rotation condition (physR). **Lower row)** Difference-ERSP (joyR minus physR), traces indicating FDR-significant (0.01) time-frequency bins.

During the outward rotation, the statistical analyses revealed distinct modulations in spectral power between the rotation conditions in the time-frequency domain. For the clusters representing neck muscle activity (Figure 3B, 1^st^ row), significant desynchronizations in the entire frequency range, most pronounced for the theta and alpha bands, were visible in the joyR condition while in the physR condition significant and long-lasting synchronizations up to 10 Hz with additional shorter bursts of synchronization in higher frequency bands were observed. For the cluster representing eye-movements (Figure 3B, 2^nd^ row), the time-warped ERSPs revealed pronounced desynchronizations in the entire frequency range during joystick rotations while no clear modulation in the lower frequency bands but significant synchronization in the alpha and higher beta bands were observed for the physical outward rotation.

Pronounced differences in time-warped ERSPs between joystick and physical rotation conditions were also visible for all brain clusters. For the RSC (Figure 3B, 3^rd^ row), power differences between physR and joyR were present in a wide frequency range from 3–60 Hz. The largest differences in terms of power increases in the RSC were observed in the 4–7 Hz theta range, as well as in the alpha and beta frequency ranges (8–30 Hz). For the parietal and occipital clusters (Figure 3B, 4^th^ and 5^th^ row, respectively), we observed desynchronization in the entire frequency range during joystick rotations following a short burst of low frequency synchronization after onset of the sphere. No such initial synchronization or later wide-band desynchronization was observed for physical rotations. In parietal cortex, both movement conditions led to an initial synchronization in the theta band with onset of the sphere. In the joyR condition, this was followed by an early desynchronization of frequencies up to 9 Hz that later extended to higher frequencies including the beta and gamma band. For physical outward rotations, an initial burst of synchronization shifted from the theta and alpha band to the alpha and beta bands. For the occipital cluster, a similar pattern of early synchronization in the theta range followed by desynchronization during outward rotation like in the parietal cluster was visible. For the physical outward rotation, in contrast only the theta range demonstrated significant desynchronizations approximately during the apex of the outward rotations accompanied by shorter initial bursts of alpha and longer lasting bursts of gamma synchronization.

## Discussion

Here, we demonstrate the modulation of neural dynamics in the human RSC and other regions of the human navigation network during heading computation in a spatial orienting task. Heading reproduction was more accurate when participants physically rotated compared to a stationary setup using a joystick, reflecting the synergistic use of idiothetic information during active movement (Klatzky et al., 1998). While an increase in heading error with increasing eccentricity of the outward rotation was observed in the joystick condition, physical rotations led to relatively low heading errors that increased only for eccentricities beyond 90 degrees pointing to general differences in the accuracy of spatial judgments based on the principal body axes (Kozhevnikov &Hegarty, 2001). The increased accuracy during physical rotations is in line with recent animal studies demonstrating improved representational accuracy for multisensory encoding of landmarks (Fischer et al., 2020; Keshavarzi et al., 2021).

Analyzing the impact of rotations on brain activity, we found the velocity of the rotation to modulate amplitudes in a wide frequency range and with topography-specific patterns. Broad-band velocity-dependent power modulations were observed for neck muscle activity only during active physical but not joystick rotations and eye movements revealed significant differences between the movement conditions in the theta and alpha frequency bands. Both the velocity-dependent modulation of neck muscle activity that was observed only during physical rotations and the differences in eye movements between the rotation conditions reflect different pursuit strategies during full body as compared to joystick-controlled rotations (Hollands et al., 2004; Robinson et al., 1986). The results thus provide a proof of principle for the analytical approach that further tested the functional relationship of rotation velocity and power modulation in different brain regions.

The RSC demonstrated velocity-dependent modulations in a broad spectrum for both physical and joystick rotations with differences between the rotation conditions in the theta and alpha frequency band. Joystick rotations revealed velocity to mainly modulate alpha band activity while physical rotations were associated with velocity-dependent modulations mainly in the theta range. These results are in line with the assumption of theta band activity underlying heading computation in the RSC integrating multisensory input. The parietal cortex, in contrast, revealed differences in power modulations between movement conditions to be restricted to the theta and alpha band mainly based on decreased grand average IC-amplitudes in the stationary setup. Further pronounced differences between the movement conditions in velocity dependent power modulations were visible in occipital cortex with modulations in the alpha frequency range in the stationary setup but no impact of velocity on amplitudes during physical rotations. These results support the assumption that occipital alpha suppression in stationary protocols reflects processing of sensory prediction errors in conditions that provide visual heading changes but no accompanying proprioceptive and vestibular information (Clark, 2013; Flossmann &Rochefort, 2021).

Further support for alpha reflecting sensory prediction error processing was revealed through time-frequency analyses during the outward rotation demonstrating significant differences in several frequency bands for brain as well as non-brain clusters. Results from neck muscle activity confirmed the prior RSA results revealing desynchronizations in a broad frequency range during joystick rotations reflecting a more rigid posture during the joystick rotation task compared to the pre-rotation baseline period. In contrast, physical rotations were accompanied by synchronized activity in the theta up to the lower beta band reflecting head rotations during the physical outward rotation. Also converging with the RSA results are pronounced differences in eye movement activity that reflected differences in pursuit eye movements for joystick-controlled (joyR) compared to pursuit eye and head movements during full body rotations (physR.) The exact nature of these differences cannot be determined without eye tracking but might be due to decreased control in the joyR condition to reduce retinal eccentricity of the moving sphere (Gouirand et al., 2019).

Notably, ERSPs over the entire time course of the outward rotation in the joyR condition displayed pronounced desynchronization of alpha oscillations in the occipital and parietal areas and, to a lesser degree, also in the RSC. Significant differences in alpha oscillations during the outward rotation are in line with similar findings of traditional desktop-based EEG studies in a variety of tasks (Klimesch, 1999), in which attention-related alpha band desynchronizations can be observed in topographically diverse areas while tasks requiring sensory-semantic information processing demonstrated alpha desynchronization with an occipito-parietal topography (Pfurtscheller &Lopes da Silva, 1999). Specifically, alpha desynchronization during heading computation was frequently documented in movement-restricted participants (Chiu et al., 2012; Gramann et al., 2010; Lin et al., 2015, 2018; Plank et al., 2010), a finding that we replicated here for the desktop-based rotation condition using a joystick and providing only visual flow information. These patterns also replicate previous results reported in Ehinger and colleagues (Ehinger et al., 2014) who used a MoBI protocol to investigate path integration. The authors investigated brain dynamics while participants were path integrating with different amounts of sensory feedback. The similarity of the results from Ehinger and colleagues’ MoBI study with previous desktop results stand in contrast to the present study potentially because the authors used a baseline in which participants actively ambulated. The present study used a baseline in which participants stood still before the rotation revealing no alpha desynchronization during active physical rotations. In contrast, pronounced broad-band increases of spectral power were present primarily in theta and beta bands. This difference in (alpha) oscillatory activity between physical and joystick rotations converges with single cell recordings during active and passive rotations in non-human primates. Here, a suppression of neural responses in the vestibular nuclei can be observed when monkeys actively rotated their head, as compared to being passively rotated on a chair (Cullen &Roy, 2004). This suppression could be due to the fact that the vestibular nuclei receive vestibular afferents as well as efferent projections from diverse sensory systems. During active head movements vestibular afferent information is suppressed by the motor-proprioceptive information generated by the active movement (Green &Angelaki, 2010). These results, together with the reported differences in velocity-dependent modulations of oscillatory activity are in line with the predictive processing approach (Clark, 2015). The frequently documented alpha desynchronization in sensory (e.g., occipital) and multimodal (e.g., parietal) cortical areas in traditional stationary EEG setups that investigate spatial orientation in movement-restricted participants might not reflect heading computation per se, but rather the processing of a significant discrepancy of predicted and perceived sensory input (Clark, 2013; Flossmann &Rochefort, 2021).

Using a MoBI approach that allows unrestricted physical movement in a spatial orientation task, we demonstrated pronounced differences in spectral modulation for heading computation based on visual flow, as compared to self-generated movements that allow utilizing idiothetic information from the visual, vestibular, and proprioceptive senses for heading computation. The results revealed a velocity-dependent modulation of brain regions that are part of the navigational network implying the relevance of velocity information from the vestibular system for heading computation. Finally, alpha desynchronization during spatial orientation tasks in traditional desktop-based setups that provide only visual flow information might point to sensory mismatch processing in cortical areas rather than heading computation itself.

## Methods

### Participants

Data were collected from 20 healthy adults (11 females) with a mean age of 30.25 years (SD = 7.68, ranging from ages 20 to 46) who received 10€/h or course credit for compensation. All participants reported normal or corrected to normal vision and no history of neurological disease. Eighteen participants reported being right-handed (two left-handed). To control for the effects of different reference frame proclivities on neural dynamics, the online version of the spatial reference frame proclivity test (RFPT^44, 45^) was administered prior to the experiment. Participants had to consistently use an ego- or allocentric reference frame in at least 80% of their responses. Of the 20 participants, nine preferentially used an egocentric reference frame, nine used an allocentric reference frame, and two used a mixed strategy. One participant (egocentric reference frame) dropped out of the experiment after the first block due to motion sickness and was removed from further data analyses. The reported results are based on the remaining 19 participants. The experimental procedures were approved by the local ethics committee (Technische Universität Berlin, Germany) and all participants signed a written informed consent in accordance with the Declaration of Helsinki. Data of the present study were further analyzed for different research purposes in Klug and Gramann (2020).

### Experimental Design and Task

Participants performed a spatial orientation task in a sparse virtual environment (WorldViz Vizard, Santa Barbara, USA) consisting of an infinite floor granulated in green and black (see figure 1B and complementary video 1). The experiment was self-paced and participants advanced the experiment by starting and ending each trial with a button press using the index finger of the dominant hand. A trial started with the onset of a red pole, which participants had to face and align with. Once the button was pressed the pole disappeared and was immediately replaced by a red sphere floating at eye level. The sphere automatically started to move around the participant along a circular trajectory at a fixed distance (30m) with one of two different velocity profiles (see Supplement for a description of the cosine functions). Participants were asked to rotate on the spot and to follow the sphere, keeping it in the center of their visual field (outward rotation). The sphere stopped unpredictably at varying eccentricity between 30° and 150° and turned blue, which indicated that participants had to rotate back to the initial heading (backward rotation). When participants had reproduced their estimated initial heading, they confirmed their heading with a button press and the red pole reappeared for reorientation. To ensure that the floor could not be used as an external landmark during the trials, it was faded out, turned randomly, and faded back in after each outward and backward rotation.

The participants completed the experimental task twice, using i) a traditional desktop 2D setup (visual flow controlled through joystick movement; “joyR”), and ii) equipped with a MoBI setup (visual flow controlled through active physical rotation with the whole body; “physR”). The condition order was balanced across participants. To ensure the comparability of both rotation conditions, participants carried the full motion capture system at all times. In the joyR condition participants stood in the dimly lit experimental hall in front of a standard TV monitor (1.5m viewing distance, HD resolution, 60Hz refresh rate, 40″ diagonal size) and were instructed to move as little as possible. They followed the sphere by tilting the joystick and were thus only able to use visual flow information to complete the task. In the physical rotation condition participants were situated in a 3D virtual reality environment using a head mounted display (HTC Vive; 2×1080×1200 resolution, 90 Hz refresh rate, 110° field of view). Participants’ movements were unconstrained, i.e., in order to follow the sphere they physically rotated on the spot, thus enabling them to use motor and kinesthetic information (i.e., vestibular input and proprioception) in addition to the visual flow for completing the task. If participants diverged from the center position as determined through motion capture of the head position, the task automatically halted and participants were asked to regain center position, indicated by a yellow floating sphere, before continuing with the task. Each movement condition was preceded by recording a three-minute baseline, during which the participants were instructed to stand still and to look straight ahead.

The starting condition (visual flow only or physical rotation) was also counterbalanced for participants with different reference frame proclivities, such that five egocentric, four allocentric, and two mixed-strategy participants started with the joyR condition, and four egocentric, five allocentric participants started with the physR condition. In each rotation condition, participants practiced the experiment in three learning trials with instructions presented on screen. Subsequently, the main experiment started, including 140 experimental trials per rotation condition. The experimental trials in each condition were randomized and split into five blocks of 28 trials each. The breaks were self-paced and the next block was initiated with the push of a button. The sphere moved either clockwise or anticlockwise around the participant; this movement was either slow or fast (randomized), depending on two different velocity profiles. The eccentricities of the sphere’s end positions were clustered from −15° to +15° around the mean eccentric end positions of 45°, 90°, and 135° in steps of 3° (e.g., the cluster 45° eccentricity ranged from 30° and 60° with 11 trials covering all eccentricities). In addition, eccentricities of 67° and 112° (2 × 8 trials) were used to achieve a near continuous distribution of end positions for the outward rotation in both rotation directions.

### Mobile Brain/Body Imaging (MoBI) setup

To allow for a meaningful interpretation of the data modalities and to preserve their temporal context, the EEG data, motion capture data from different sources, and experiment event marker data were time-stamped, streamed, recorded, and synchronized using the Lab Streaming Layer (Kothe, 2014).

### Data Recordings: EEG

EEG data was recorded from 157 active electrodes with a sampling rate of 1000 Hz and band-pass filtered from 0.016 Hz to 500 Hz (BrainAmp Move System, Brain Products, Gilching, Germany). Using an elastic cap with an equidistant design (EASYCAP, Herrsching, Germany), 129 electrodes were placed on the scalp, and 28 electrodes were placed around the neck using a custom neckband (EASYCAP, Herrsching, Germany) in order to record neck muscle activity. Data were referenced to an electrode located closest to the standard position FCz. Impedances were kept below 10kΩ for standard locations on the scalp, and below 50kΩ for the neckband. Electrode locations were digitized using an optical tracking system (Polaris Vicra, NDI, Waterloo, ON, Canada).

### Data Recordings: Motion Capture

Two different motion capture data sources were used: 19 red active light-emitting diodes (LEDs) were captured using 31 cameras of the Impulse X2 System (PhaseSpace Inc., San Leandro, CA, USA) with a sampling rate of 90 Hz. They were placed on the feet (2 × 4 LEDs), around the hips (5 LEDs), on the shoulders (4 LEDs), and on the HTC Vive (2 LEDs; to account for an offset in yaw angle between the PhaseSpace and the HTC Vive tracking). Except for the two LEDs on the HTC Vive, they were subsequently grouped together to form rigid body parts of feet, hip, and shoulders, enabling tracking with six degrees of freedom (*x, y*, and *z* position and *roll, yaw*, and *pitch* orientation) per body part. Head motion capture data (position and orientation) was acquired using the HTC Lighthouse tracking system with 90Hz sampling rate, since it was also used for the positional tracking of the virtual reality view. Because the main focus of the study concerned the head movement-related modulation of neural dynamics in RSC, only data streams from the head motion capture data were used for the analysis.

### Data Analysis

Data analysis was done in MATLAB (R2016b version 9.1; The MathWorks Inc., Natick, Massachusetts, USA), using custom scripts based on the EEGLAB (Delorme &Makeig, 2004), MoBILAB (Ojeda et al., 2014), RSA (Nili et al., 2014), and in-house toolboxes (Klug &Gramann, 2020).

### Motion Capture Data Analysis: Automatic Detection of Movement Markers

Motion capture data was preprocessed using MoBILAB (Ojeda et al., 2014) with adapted functions. The rigid body data was recorded in x, y, z, as well as quaternion orientation values, a 6 Hz zero-lag lowpass FIR filter was applied to the data, the orientation was then transformed into Euler angles and three time derivatives were subsequently calculated. For movement detection the absolute velocity of head orientation (yaw) was used (“physR” condition: visual flow controlled by head MoCap; “joyR”: visual flow controlled by the joystick). For convenience, in the remainder the term “head movement” refers to both rotation conditions. Head movement onset and offset events were extracted based on the velocity: A movement onset was initially defined as having a greater velocity than the 65% quantile of the complete data set (estimating movement to happen 35% of the time during the experiment). Once this coarse threshold was reached, the movement onset and offset event markers were created based on a finer threshold of 5% of the maximum velocity in a window of 2s around the detected movement. New movements could only be detected after the offset of the previous movement. A minimal movement duration of 285ms was defined to exclude movement artefacts created by jitter in the MoCap recording. The final data was exported as EEGLAB data set, synchronized, and different streams (EEG, MoCAP) were split into separate sets to allow for EEG-specific analysis based on movement markers.

### Behavioral Data Analysis – Heading Error

#### Absolute heading error

For each epoch the absolute heading error was defined by taking the absolute difference between the participant’s initial start orientation (yaw; Euler angles) and the participant’s orientation after completing the backward rotation (completion indicated by the button press). The absolute heading error gives a robust overall indication of deviations from the starting orientation, without considering direction-specific over-or underestimation. For each participant and condition occasional outlier epochs with errors larger than three standard deviations were excluded. Furthermore, filler trials were also excluded. Finally, the absolute heading errors from the remaining valid epochs were averaged for three eccentricity categories within a range of ± 15° (45°: 30–60°; 90°: 75–105°; 135°: 120–150°). The term “heading error” refers to the average within each eccentricity category.

#### Statistics

The group-level statistics were performed with SPSS (version 25; IBM SPSS Statistics for Windows, Armonk, NY: IBM Corp.). For the absolute heading error a 2 × 2 × 3 factorial repeated measures analysis of variance (rANOVA) was performed with the within-participant factors “rotation condition” (physR, joyR), “direction” (clockwise, anti-clockwise), and “eccentricity” (15°, 30°, 45°); for the signed heading error cf. Supplements. In case the assumption of sphericity was violated, Greenhouse-Geisser corrected values are reported. If required, post hoc analysis was performed with the paired t-test. If the majority of data sets (≥ 50%) were not normally distributed (Kolmogorov-Smirnov test), post hoc testing was performed with the paired Wilcoxon signed-rank test. Bonferroni correction for post hoc testing was used in the case of multiple comparisons. Raw p-values are reported and indicated as significant if they were lower than the Bonferroni adjustment significance threshold. In general, while being aware that non-parametric post hoc testing was performed on ranks, for convenience the average values are presented as mean ± standard deviation (SD).

### Independent Component Analysis (ICA)

Artefactual channels were manually identified, removed, and interpolated using spherical interpolation (on average, 17.6 channels were interpolated, SD=9.5). Subsequently, the data was re-referenced to the average of all channels and a zero-phase Hamming windowed high-pass FIR filter (order 827, pass-band edge 1 Hz) was applied to the data. Then artefactual time segments in the data were manually rejected; eye movements were not considered as artefacts. Data of both rotation conditions and their respective baselines were appended, and subsequently, the data was parsed into maximally independent components (IC), using an adaptive mixture independent component analysis (AMICA) algorithm (Palmer et al., 2006) with a principal component analysis (PCA) reduction to the remaining rank of the data set. For each IC an equivalent dipole model was computed as implemented by DIPFIT routines (Oostenveld &Oostendorp, 2002). For this purpose the individually measured electrode locations were rotated and rescaled to fit a boundary element head model (BEM) based on the MNI brain (Montreal Neurological Institute, MNI, Montreal, QC, Canada). We refer to the approximated spatial origin of an IC as “in or near” the specified location.

### Event-Related Spectral Perturbations (ERSP)

A new copy of the pre-processed EEG data set was created, comprising interpolated channels and the complete continuous time courses, and the computed IC spatial filters and their respective source localization estimates were copied to this data set. Epochs of 13s length were created around the sphere onset markers, including a 1s pre-stimulus interval, resulting in 140 epochs per participant and condition. A spectrogram of all single trials was computed for all IC activation time courses using the newtimef() function of EEGLAB (3 to 100 Hz in logarithmic scale, using a wavelet transformation with 3 cycles for the lowest frequency and a linear increase with frequency of 0.5 cycles). To compute the final ERSP for the RSC region, artefactual epochs were first automatically rejected based on MoCap data and the IC activation time courses present in the respective cluster by removing epochs with baselines contaminated by head movements and epochs that contained considerable artefacts in the IC activation time course during task performance (as evaluated by epoch mean, standard deviation, and Mahalanobis distance (Nierula et al., 2013; see Supplements for more details). Artefact cleaning resulted in a sum of 1566 ERSP epochs for the physical rotation condition and 1880 epochs for the joystick rotation condition.

### EEG Data Group-Level Analysis

#### Repetitive clustering approach

To allow for a group-level comparison of EEG data at the source level (ICs), the 70 ICs of each participant explaining most of the variance of the data were selected (1330 ICs in total) and subsequently clustered based on their equivalent dipole locations (weight=6), grand-average ERSPs (weight=3), mean log spectra (weight=1), and scalp topography (weight=1), using a region of interest (ROI) driven repetitive *k*-means clustering approach. The weighted IC measures were summed and compressed using PCA, resulting in a 10-dimensional feature vector for clustering. ICs were clustered by applying the *k*-means algorithm with n=50 cluster centroids to the resulting vectors and their respective distance between each other in vector space. We chose to use fewer clusters than ICs per participant because of our assumption that, although statistically independent per time point, there may be more than one IC per participant that is similar in function and location. ICs with a distance of more than three standard deviations from any final centroid mean were considered outliers.

Crucially, to ensure replicability of the clustering, we clustered 10,000 times and selected the final solution based on the following approach: i) We defined the Retrosplenial Complex (RSC) as the ROI in Talairach coordinates (*x*=0, *y*=−45, *z*=10) for which the clustering should be optimized; ii) for each resulting cluster in the region of interest (cluster of interest; COI), we calculated the number of participants with an IC in the cluster, the ratio of the number of ICs per participant in the cluster, the spread of the cluster (average squared distance of each IC from the cluster centroid), the mean residual variance (RV) of the fitted dipoles in the cluster, x, y, z coordinates of the cluster centroid, the distance of the cluster centroid from the ROI, and the Mahalanobis distance of this COI from the median of the distribution of the 10,000 solutions. The x, y, z coordinates were only used to determine the Mahalanobis distance and were not in the final quality measure vector; iii) the quality measures were standardized to their respective maximum, then weighted (#participants: 2, ICs/participants: −3, spread: −1, RV: −1, distance from ROI: −3, Mahalanobis distance from the median: −1). These weights optimize for a clustering solution that contains a cluster close to the ROI of interest, which contains the ICs of many participants, but a low number of ICs per participant. Finally, the solutions were ranked according to their summed score, and the highest ranked solution was chosen as the final clustering solution. The following “brain” and “non-brain” clusters were selected for further analyses, containing at least 70% of participants.

#### Clustering results for RSC (cluster #16)

The final solution for the RSC cluster contained the ICs from 15 participants, a ratio of 1.13 ICs per participant, a mean RV of 10.3%, and a spread of 296 with a distance of 10.2 units in the Talairach space (both measures with regards to the ROI coordinates).

The final solution for the right parietal cluster contained 20 ICs from 14 participants, a ratio of 1.43 ICs per participant, and a mean RV of 12.4%. The final solution for the occipital cluster contained 22 ICs from 15 participants, a ratio of 1.46 ICs per participant, and a mean RV of 8.2 %. For the eye cluster, 26 ICs from 14 participants were included with a ratio of 1.86 ICs per participant and a mean RV of 23.1%. The neck cluster contained 20 ICs from 16 participants, a ratio of 1.25 ICs per participant and a mean RV of 34.8 %.

#### Cluster cleaning with IClabel

Each selected cluster was further automatically analysed for potential “artifact” ICs using ICLabel (Pion-Tonachini et al., 2019). This algorithm allows to classify ICs into “brain” or “non-brain” origin, which was previously used in the context of evaluating different pre-processing approaches for mobile EEG recordings (same data set of the present study; Klug &Gramann, 2020). IClabel was run in the “default” version (beta version, 2017), using IC topography, autocorrelation, and spectrum for the classification and the selected classes were “brain” and “other” with a threshold of 0.7. After the classification n=15 participants remained for the RSC cluster, n=14 for parietal, n=15 for occipital, n=14 for the eye cluster and n=16 for the neck cluster, respectively.

#### ERSP group-level analysis

ERSPs were computed for the RSC cluster by first averaging the time-frequency data at the IC level, then at the participant level, and finally at the group level. The time-frequency data of each trial was normalized by its mean activity (Grandchamp &Delorme, 2011) and the average ERSP for each IC was calculated and baseline-corrected using a divisive baseline (mean activity in the interval of 200ms prior to sphere movement onset). Subsequently, the ERSPs of all ICs per participant were averaged. Finally, the ERSPs of all participants were averaged and log-transformed from the power-space into decibels (dB=10*log10(power)). Statistical analysis comparing ERSP activity either against the baseline or between conditions was performed at the group level using a permutation test with 2,000 permutations, and a multiple comparison correction using the false discovery rate (FDR; α=0.01). Final plots contain the significance thresholds as contours of significant time-frequency bins.

### EEG activity associated with varying head movement velocity

In order to estimate differential EEG activation depending on head movement velocity (slowest to largest) and frequency band (4-30.5 Hz), data of each IC (Hilbert-transformed (Clochon et al., 1996); baseline-corrected amplitude during the outward rotation epochs, i.e., movement onset to offset), baseline-corrected amplitude during the outward rotation epochs, i.e., movement onset to offset) was subjected to a velocity binning procedure (Bassett &Taube, 2001; Linkenkaer-Hansen et al., 2004) and Mahalanobis distance-based Representational Similarity Analysis (Kriegeskorte &Kievit, 2013; Nili et al., 2014; Tanaka et al., 2018). For more details on the velocity binning, RSA (including single participant examples), and statistics please refer to the Supplements.

## Acknowledgements

The research was supported by the German Research Foundation (DFG) grant no. GR 2627/8-1. We gratefully acknowledge the help of Jonna Jürs and Yiru Chen with the data acquisition and pre-processing.

## Author contributions

K.G. conceived the study and obtained funding, K.G. designed the protocol, M.K. and L.G. performed recordings, F.H., M.K., and L.G. analysed data. All authors interpreted data and discussed results. K.G. and F.H. wrote the manuscript. All authors commented and edited the manuscript.

## Additional information

The authors report no conflict of interest

