## Supplement for "Human cortical dynamics during full-body heading changes"

**Supplements**

**Supplements – Video material**

*Video ‘Spot_Rotation_2018_joyR.mp4’*. The video displays a participant in the joyR condition positioned in front of the screen and following the sphere during a rotation to the left via joystick control. One trial is commented according to the different phases of the experiment. The video is available at: <https://osf.io/qrw9d/>

*Video ‘Spot_Rotation_2018_physR.mp4’*. The video displays a participant in the physR condition in the MoBI setup with HMD including a rigid body for motion capture and EEG, following the sphere during a rotation to the left via physical rotation with the whole body. One trial is commented according to the different phases of the experiment. The video is available at: <https://osf.io/6mfzg>

**Supplements – Methods I: Experimental design and task**

*Velocity Profiles*. Upon button press, a red sphere started to travel at 5 m/s along a circle with a 30 m radius and accelerated either to a maximum of 35 m/s (fast condition) or to a maximum of 30 m/s (slow) condition. The acceleration and deceleration profiles were stretched or compressed to the eccentricity of the current trial. Eccentricity indicates the angle away from the participant when facing the initial local landmark (pole).

### Supplements – Methods II: Behavioural data

*Relative heading error*. When considering relative differences, i.e., signed errors, for instance for clockwise rotations a negative error refers to an “undershoot“ (i.e., the participant did not fully rotate back to the initial start orientation), whereas a positive error refers to an “overshoot“ (i.e., the participant rotated further than the initial start orientation). Notably, for anti-clockwise rotations the sign for over- and undershoot is reversed. However, in order to have matching signs for both rotation directions, the relative error values were inverted for the anti-clockwise rotation epochs before averaging across valid epochs. Previously identified occasional outlier and filler epochs were excluded (cf. absolute heading error), and the remaining valid epochs were averaged. The group level statistics were performed as the absolute heading error.

*Reaction times*. The reaction time of the outward rotation was determined for each available epoch, defined as the time difference between the appearance of the visual stimulus (i.e., red sphere) and the first subsequent movement onset (cf. automatic detection of movement markers, as described in the Methods). In order to exclude epochs that contained excessively strong movements, which could have impeded an accurate detection of the movement onset, epochs were defined as invalid and were rejected if at least one of the following criteria was violated: i) if the yaw range exceeded 5° in the pre-stimulus interval (-500 ms to stimulus onset); ii) if the yaw range exceeded 5° in the pre-movement interval (stimulus onset to movement onset); iii) the first detected movement marker after the visual stimulus was an offset, not an onset; iv) no detected movement onset within the outward rotation epoch (i.e., appearance until disappearance of the visual stimulus). For each participant and condition, the reaction times were averaged across the remaining valid epochs, subsequently referred to by the term “reaction time“. A 2 x 2 factorial rANOVA was performed with the within-participant factors “rotation condition“ (physR, joyR) and “direction“ (clockwise, anti-clockwise). For further details on the group level statistics cf. the absolute heading error section.

*The Simulator Sickness Questionnaire (SSQ)*. The Simulator Sickness Questionnaire (Kennedy et al., 1993) was administered to each participant three times: prior to any experimental recordings (“baseline“) and after each of the experimental conditions (physR, joyR). For each participant the mean value across all items (Likert scale, ranging 0-4) was calculated separately for each SSQ subscore (“nausea“, “oculomotor“, “disorientation“ score). Subsequently, the term “SSQ“ refers to the mean across items. Separately for each “SSQ“ subscore (“nausea“, “oculomotor“, “disorientation“ score) a oneway rANOVA was performed with the within-participant factor “rotation condition“ (physR, joyR). For further details on the group level statistics cf. the absolute heading error section.

*The Igroup Presence Questionnaire (IPQ).* The Igroup Presence Questionnaire (Schubert et al., 2001) was administered to each participant once after performing the physR condition. For each participant the mean value across all items (Likert scale, ranging 0-6) was calculated separately for each IPQ subscore (“spatial presence“, “involvement“, “experienced realism“), whereas “general presence“ consisted of one item only.

#### Supplements – Methods III: Automatic cleaning of ERSP epochs

For each rotation condition the ERSP epochs were automatically cleaned by three approaches: i) in order to exclude possible contamination of the ERSP baseline, epochs that contained strong head movements (i.e., orientation yaw exceeding 5°) in the pre-stimulus interval and/or pre-movement interval were excluded for baseline calculation; ii) epochs that contained large head movement before the movement onset event (exceeding 5°; indicating a miss of the automatic movement onset detection) and/or the first detected movement marker after the visual stimulus was an offset, not an onset; iii) 10 % of epochs were automatically removed ranking largest for contamination by artefacts, as described below.

*Automatic epoch removal by ranking*. For iii) the epochs (absolute values of IC time courses) were separately ranked with respect to their maximum value, as determined by three approaches: “standard deviation”, “Mahalanobis distance”, and “mean”. First, the time course in each epoch was averaged for each selected IC, denoted as *averageIC_epoch_*. Then each epoch was evaluated separately by each of the three approaches:

A) “Standard deviation“. The standard deviation was calculated for *averageIC_epoch_*, resulting in a single value per epoch. Increasing values indicate more diverse IC activity in the epoch, thus being a sensitive measure for the detection of several “outlier“ IC (or channels, if performed in sensor space).

B) “Mahalanobis distance”. The Mahalanobis distance (Mahalanobis, 1936) was calculated for *averageIC_epoch_*, resulting in a single value per epoch (Nierula et al., 2013). Complementary to A), it considers also the covariance between IC/channels, and in practice the Mahalanobis distance appeared to be sensitive especially to the presence of only few yet strong outliers.

C) “Mean”. The mean across *averageIC_epoch_* was calculated, obtaining a single value per epoch (i.e., averaged across time and all IC). This measure represents an indicator of contaminations occurring in many IC/channels, e.g., due to strong baseline shifts or technical artefacts.

*Ranking procedure*. After obtaining numeric indices for each epoch separately by the three approaches, for each approach the indices were ranked in ascending order. Furthermore, in order to take into account the possibility that different epochs were detected by the three approaches (and thus possibly leading to unintended data loss more than the initially threshold, here 10%), the indices were weighted separately (i.e., ranks multiplied by the weighting factor [3 1 1] for “Mahalanobis distance”, “standard deviation”, and “mean”, respectively). Finally, the sum across weighted ranks was calculated and sorted in ascending order. Based on this final epoch rank list the 10% “worst“ epochs were removed. This approach takes all three approaches into account, thus allowing sensitivity for different kinds of artifacts, while at the same time avoiding unintended data loss.

The final set comprised 127.5 ± 15.8 (mean ± SD) epochs for the baseline in the physR condition and 131.9 ± 16.1 in the joyR condition. On average 104.4 ± 12.6 epochs (physR) and 125.0 ± 9.6 epochs (joyR) were considered for the ERSP data per rotation condition, resulting in a total sum of 1566 and 1880 epochs for the physR and joyR condition, respectively.

#### Supplements – Methods IV: EEG activity associated with varying head movement velocity

*Data preprocessing.* One interest of the present study was the modulation of oscillatory brain activity associated with velocity changes (orientation; yaw) of visual flow (joyR condition; visual flow controlled by joystick movement) as well as for actual head movements (motion capture in the physR condition: visual flow controlled by head orientation), both referred to by “head movement velocity“. For this purpose the analysis was based on the Mahalanobis distance (cf. Representational Similarity Analysis [RSA]; Kriegeskorte & Kievit, 2013; Nili et al., 2014) of EEG amplitudes from single-trial velocity binning, as described below.

Three “brain activity” clusters (RSC, right parietal cortex, occipital cortex) and, for comparison purposes, two “non-brain activity” clusters (vertical eye movements and left-sided neck muscular activity) were included in the subsequent analysis. Since not all participants were equally represented in all clusters, the analysis was performed for each cluster separately. For each component in a given cluster, amplitude envelopes (Hilbert transform; Clochon et al., 1996) of band-pass filtered (Butterworth filter, fourth order) continuous oscillatory activity in nine major frequency bands (non-overlapping, step 2.5 Hz) were obtained in the range from 4-30.5 Hz (4-6.5, 7-9.5, 10-12.5, 13-15.5, 16-18.5, 19-21.5, 22-24.5, 25-27.5, and 28-30.5 Hz). The frequency bands in theta, alpha, and beta ranges were selected with respect to previous findings suggesting their involvement in task-relevant domains, such as spatial navigation (Chiu et al., 2012; Do et al., 2020; Gramann et al., 2010; Lin et al., 2015, 2018; Plank et al., 2010) and sensorimotor processing (Engel & Fries, 2010; Pfurtscheller et al., 1997). EEG data were shifted back by 100 ms in order to match the MoCap recordings (due to technical acquisition delays of the multivariate data streams.

In each frequency band for each IC in a given cluster, paired EEG and MoCap velocity epochs were created with respect to visual stimulus onset (i.e., start of the outward rotation): “baseline“ (B_EEG_) referring to -200 ms to stimulus onset, and “movement“ (M_EEG_, M_velocity_) epochs, referring to the first movement onset to offset after the stimulus, according to the automatically determined velocity onset and offset marker in the continuous MoCap data stream (head orientation; yaw) as described in the methods section of the manuscript. In order to ensure sufficient data quality, M_velocity_ epochs were excluded that were too short (< 500 ms) and/or too long (> 15 s). Furthermore, B_velocity_ epochs were excluded that might have been contaminated by occasional movements, i.e., if containing movement onset or offset markers (automatic detection, see above) and if the orientation yaw was > 5°. All “invalid” epochs were deleted from all data sets (B_EEG,_ B_velocity_, M_EEG_, M_velocity_). Subsequently, additional rigid EEG artefact rejection was applied, removing 20 % of M_EEG_ epochs with the largest mean value (sorted in ascending order) from all data sets. Afterwards M_EEG_ epochs from the remaining “valid” epoch pool were baseline-corrected (epoch-wise) by subtracting the mean value across the baseline interval in the respective epoch: EEG_diff_ = M_EEG_ – mean(B_EEG_).

*Single trial movement velocity binning analysis.* In order to assess modulation of EEG oscillations accompanying different movement velocities, a binning procedure was applied (Bassett & Taube, 2001; Linkenkaer-Hansen et al., 2004) for each epoch separately in each rotation condition: In a given M_velocity_ epoch percentiles (10 percent steps; resulting in 10 bins) were taken of all velocity values (sorted in ascending order). Subsequently, all the data samples were assembled that corresponded to each of the percentile categories. Notably, this “relative binning“ procedure by taking percentiles (in contrast to the “absolute binning“ of previous work, i.e., equidistant velocity binning (Bassett & Taube, 2001) has the advantage of providing equal amounts of samples per velocity percentile bin. Therefore, this approach avoids a considerable drawback of absolute velocity binning which would provide only very few data samples for higher (i.e., less frequent) velocity values.

After having obtained the velocity bins for the given epoch, EEG_diff_ was averaged across all data samples in a given percentile bin, obtaining EEG_diffMeanPerBin_. Finally, the single-epoch EEG_diffMeanPerBin_ matrix was averaged across all valid epochs, obtaining EEG_diffMeanPerBinAv_.

This procedure was repeated for all epochs of a given IC in a given clusters, separately for each frequency band and separately in each rotation condition. If occasionally multiple IC were available per participant in a given cluster, the obtained EEG_diffMeanPerBinAv_ values were averaged across the available IC for the respective participant, thus obtaining a single value per participant and velocity bin, obtaining EEG_diffMeanPerBinAvIC_. These finally obtained EEG_diffMeanPerBinAvIC_ values basically represent average changes in EEG (IC) amplitude with respect to the pre-stimulus baseline for slow to fast movement velocity ranges (as defined by the percentile categories) estimated across all valid trials.

*Representational similarity analysis (RSA).* The RSA framework (Kriegeskorte & Kievit, 2013; Nili et al., 2014) provides a powerful approach to visualize and quantify similarities/distances between data obtained from multiple measures and/or in multiple conditions. This framework was selected in the present study, given that for a single participant in a given cluster a 2 x 90 (= 180 values) data matrix (based on EEG_diffMeanPerBinAvIC_ ) was available (two rotation conditions, 9 frequency bands, 10 velocity bins; containing the baseline-corrected, averaged EEG amplitudes). Despite from its frequent application in fMRI (Kriegeskorte & Kievit, 2013), RSA was recently utilized in the EEG context for visual processing (Groen et al., 2012) and for evaluating directional tuning of neural activity by multiple movement directions (finger pointing; Tanaka et al., 2018). In the present study the RSA framework (Kriegeskorte & Kievit, 2013) was utilized to evaluate neural activation across multiple velocities, frequency bands, and rotation conditions based on their “representational geometry”, i.e., at the level of similarity (here: estimated by the Mahalanobis distance) rather than based on differences of actual EEG amplitudes. For this purpose, a “representational dissimilarity matrix” (RDM; Kriegeskorte & Kievit, 2013) was derived for each participant based on the EEG_diffMeanPerBinAvIC_ matrix (180 values) by calculating the rank-transformed normalized Mahalanobis distance for each field, as derived from the RSA toolbox (Nili et al., 2014), denoted as MD_norm_, which is scaled from 0 to 1 (max.) distance. Subsequently, the more general term RDM will refer to the MD_norm_ values (180 x 180 values), also denoted here as RDM_observed_.

Additionally, for statistical comparison purposes RDM_shuffled_ was calculated for each participant in a given cluster: the EEG_diffMeanPerBinAvIC_ data matrix was randomly shuffled, thus destroying systematic relations between velocity bins and EEG amplitudes, and RDM_shuffled_ was calculated on the shuffled data matrix. This was repeated for n = 1000 rounds and subsequently all the obtained RDM_shuffled_ were averaged across all rounds, constituting RDM_shuffledAV_.

Finally, in a given cluster the respective RDM of all participants were assembled (RDM_observed_all_, RDM_shuffledAV_all_).

*Statistics*. One main approach in the RSA framework is the comparison between “observed” RDM vs. “reference” RDM, the latter being based on pre-selected computational or theoretical models (Nili et al., 2014) of well-known neural phenomena such as face/house distinction in the fusiform face area (Kriegeskorte & Kievit, 2013). However, in the present study no such a priori models were available, since this study investigated for the first time movement velocity-associated modulation of neural activity during spatial orientation in actively moving humans. Therefore, two approaches were selected in order to test for the presence of “significant” MD_norm_ values across all participant in a given cluster: i) testing vs. 0 (as demonstrated in the RSA toolbox; Nili et al., 2014), and ii) permutation testing (Pesarin & Salmaso, 2010) of MD_norm_ being larger than its noise estimate.

*Statistics I: testing vs. 0*. The first approach was utilized in order to test for the presence of “significant” MD_norm_ values in general, i.e., being larger than 0. Separately in each cluster, each matrix field (row *r*, column *c*) of RDM_observed_all_ (i.e., containing RDMs from all available participants) was compared to 0 (one sample t-test, right-tailed). Since the Mahalanobis distance even of random values would be comparatively small but not zero, this leads to an avoidable positive bias when testing vs. 0. Therefore, a strict correction for multiple comparisons with the False Discovery Rate (FDR; Benjamini & Yekutieli, 2001) at p = 0.0001 level was applied. Non-significant results were masked.

*Statistics II: permutation testing*. The second, more conservative approach of non-parametric permutation testing (Pesarin & Salmaso, 2010) was utilized in order to test for the presence of “significant” MD_norm_ vs. its noise estimate (MD_norm_ calculated on shuffled EEG data), i.e., by obtaining in each matrix field (row *r*, column *c*) the error probability (right-tailed; threshold p = 0.05) for the sample means RDM_observed_all_ being larger than RDM_shuffledAV_all_ across n = 10000 permutations. Non-significant results were masked.

All Supplementary results are presented as (mean ± SD).

### Supplements - Results

**Heading error**

The analysis of heading errors comprised 131 ± 1 trials in both, the physR and joyR condition, respectively.

*Relative heading error*. The rANOVA revealed a significant main effect of “direction“ (F_1,18_ = 6.82; p = 0.018) and of “eccentricity“ (F_1.24,22.34_ = 33.63; p < 0.001). Two significant interaction effects were present: i) between “rotation condition“ and “eccentricity“ (F_1.58,28.42_ = 13.27; p < 0.001) and ii) between “rotation condition“, “eccentricity“, and “direction“ (F_2,36_ = 3.54; p = 0.039). Post hoc analysis (paired t-test) of the latter interaction effect ii) revealed that for both rotation directions the smallest eccentricity category (45°) was associated with a positive error (i.e., overshoot, too far rotation), while the largest eccentricity category (135°) was associated with a negative error (i.e., undershoot, too short rotation). Furthermore, direction-specific differences between both rotation conditions were obtained. Summarizing the results: i) in both rotation conditions smaller eccentricities (~ 45°) were associated rather with error overshoots (on average 6.73°), i.e., the back rotation exceeded the initial start orientation, whereas larger eccentricities (~ 135°) were associated rather with error undershoots (on average -6.67°), i.e., the back rotation was too short and the participants did not reach the initial start orientation; ii) the relative heading error was significantly larger for the joyR condition than for the physR condition, depending on the rotation direction and the eccentricity (anti-clockwise: 45°; clockwise: 135 °), which in actual terms suggests that for small/large eccentricities in the joyR condition the participant had a tendency to stop the back rotation not at about 0°, but rather showing a slight shift to the left stopping at about -12°. The statistical results are presented in Supplements Fig. 1. While the analysis of more differentiated eccentricity categories was beyond the scope of the present study due to a limited number of trials, visual inspection of the relative heading errors in decimal steps revealed for both conditions and rotation directions a flip from “overshoot“ to “undershoot“ errors at about 110° (cf. Supplements Fig. 2A-B).

#### Reaction times

*Outward rotation*. The reaction time comprised 110 ± 20 (mean ± SD) and 127 ± 20 epochs in the physR and joyR condition, respectively. The rANOVA revealed a significant main effect of “rotation condition“ (F_1,18_ = 68.37; p < 0.001), showing significantly shorter reaction time for “physR“ (0.48 ± 0.22 s) than for “joyR“ (1.12 ± 0.31 s). No further significant effects were obtained. Results are presented in Supplements Fig. 3.

#### Simulator Sickness Questionnaire (SSQ) & Igroup Presence Questionnaire (IPQ)

*IPG*. Overall, the IPG indicated that the participants experienced a relatively degree of presence during the physR condition (general presence: 3.63 ± 1.71; spatial presence: 4.31 ± 1.36; involvement: 2.75 ± 1.56; experienced realism: 1.55 ± 1.06).

*SSQ*. In general, while overall the SSQ scores were very low (grand-averages < 1), SSQ in both rotation conditions was slightly increased compared to the baseline. However, SSQ for both rotation conditions was similar in each of the subscores (mean ± SD):

*SSQ - Nausea subscore*. The rANOVA revealed a significant main effect of “rotation condition“ (F_1.48,26.57_ = 7.93; p = 0.004). Post hoc analysis (Wilcoxon signed rank test) revealed slight albeit significant differences, with “baseline“ (0.06 ± 0.1) being smaller than “physR“ (0.26 ± 0.2; p = 0.002) and than “joyR“ (0.3 ± 0.34; p = 0.004), respectively. No further significant effects were obtained.

*SSQ - Oculomotor subscore*. The rANOVA revealed a significant main effect of “rotation condition“ (F_2,36_ = 10.44; p < 0.001). Post hoc analysis (Wilcoxon signed rank test) revealed slight albeit significant differences, with “baseline“ (0.23 ± 0.21) being smaller than “physR“ (0.61 ± 0.49; p = 0.002) and than “joyR“ (0.53 ± 0.47; p = 0.006), respectively. No further significant effects were obtained.

*SSQ - Disorientation subscore*. The rANOVA revealed a significant main effect of “rotation condition“ (F_2,36_ = 7.45; p = 0.002). Post hoc analysis (Wilcoxon signed rank test) revealed slight albeit significant differences, with “baseline“ (0.05 ± 0.07) being smaller than “physR“ (0.27 ± 0.27; p = 0.003), but no Bonferroni-significant differences to “joyR“ (0.14 ± 0.27; p = 0.03). No further significant effects were obtained.

#### EEG differences associated with varying head movement velocity

*Single trial movement velocity binning*. On average artefact-cleaned 109 ± 3 “movement“ epochs (physR: 108 ± 4; joyR: 111 ± 2) were subjected to the velocity binning analysis, i.e., referring to the “movement“ interval of the outward rotation (as defined by velocity onset to offset after the visual stimulus; head orientation yaw). The average “movement“ epoch duration was 3.9 ± 0.7 s (physR: 3.8 ± 0.7 s; joyR: 4 ± 0.7 s). The ranges for the obtained velocity bins were on average (percentile bin 1-10: ≥ start until < end): 0-9°/s (± 1); 9-13°/s (± 2), 13-16°/s (± 3), 16-19°/s (± 4), 19-22°/s (± 4), 22-26°/s (± 5), 26-30°/s (± 5), 30-36°/s (± 4), 36-41°/s (± 5), 41-47°/s (± 5). The average duration per velocity bin was 400 ms ± 100 ms.

*Representational similarity analysis (RSA) – single participants.* Examples of representative single participants are presented in Supplements Fig. 4, showing i) the baseline-corrected, trial-averaged amplitude of component (IC) activity per frequency band and movement velocity bin which was subjected to RSA, ii) the resulting RDM calculated for the given IC activity (RDM_observed_), and iii) the RDM on the shuffled IC activity (RDM_shuffledAV_). As expected, no pronounced patterning of distances was observed for the shuffled data (MD_norm_ of ~ 0.5; fourth column), whereas distinct patterning in the MD_norm_ values was observed in the original data sets (third column). Notably, varying movement velocity was selectively associated with differential IC activity depending on the topography (brain vs. non-brain IC), frequency band, and rotation condition, as reflected in pronounced “striping” in the IC amplitudes and in the RDM. Furthermore, the RDM results clearly indicate an advantage of using the (normalized) Mahalanobis distance measure to evaluate similarities in neural activity (instead of actual amplitude differences), since the MD_norm_ is scale-invariant to the actual amplitude; for instance, increasing movement velocity in the physR condition is associated with a relative increase of IC amplitude in the neck (approx. 13-27 Hz) and with a relative amplitude decrease in the right parietal cortex (approx. 7-18 Hz), whereas both reactivity patterns are associated with increasing distance in the RDM (vertical striping) irrespective of actual amplitudes, which might vary according to the topography or presence of noise.

*RSA – group level analysis.* Results were obtained for two “non-brain” clusters (participants: eye n = 14, right neck n = 16) and for three “brain” clusters (participants: RSC n = 15, right parietal n = 14, occipital n = 15). On average, the noise estimate of MD_norm_, i.e., distances calculated on the shuffled IC data (RDM_shuffledAV_), was approx. 0.5 ± 0.002 (mean ± SD) across all presented clusters, with average min. values of 0.49 and average max. values of 0.51. Subsequently, results of statistical testing are presented for each comparison type: physR (4005 fields of the RDM, i.e., velocity bin x band combinations excluding the main diagonal), joyR (4005 fields), and physR vs. joyR (8100 fields).

*RSA – group level statistics for cluster: right neck.* Grand-average RDM and maps of significant p-values are shown in Figure 2. I) testing vs. 0: When testing for “presence” of distances per se, FDR-significant results were obtained for 92 % of RDM fields in the physR condition (upper left quadrant), for 28 % in the joyR condition (lower right quadrant), and for 94 % in the physR vs. joyR comparison (lower left quadrant). II) testing vs. shuffled: In the more conservative approach, i.e., testing differences against their noise estimate, permutation-significant results were obtained for 39 % (physR) of RDM fields, 0 % (joyR), and 30 % (physR vs. joyR).

*RSA – group level statistics for cluster: eye.* Grand-average RDM and maps of significant p-values are shown in Figure 2. I) testing vs. 0: FDR-significant results were obtained for 51 % (physR) of RDM fields, 68 % (joyR), and 82 % (physR vs. joyR). II) testing vs. shuffled (noise estimate): Permutation-significant results were obtained for 20 % (physR) of RDM fields, 35 % (joyR), and 40 % (physR vs. joyR).

*RSA – group level statistics for cluster: RSC.* Grand-average RDM and maps of significant p-values are shown in Figure 2. I) testing vs. 0: FDR-significant results were obtained for 52 % (physR) of RDM fields, 62 % (joyR), and 65 % (physR vs. joyR). II) testing vs. shuffled (noise estimate): Permutation-significant results were obtained for 7 % (physR) of RDM fields, 14 % (joyR), and 15 % (physR vs. joyR).

*RSA – group level statistics for cluster: right parietal cortex.* Grand-average RDM and maps of significant p-values are shown in Figure 2. I) testing vs. 0: FDR-significant results were obtained for 33 % (physR) of RDM fields, 44 % (joyR), and 67 % (physR vs. joyR). II) testing vs. shuffled (noise estimate): Permutation-significant results were obtained for 5 % (physR) of RDM fields, 16 % (joyR), and 33 % (physR vs. joyR).

*RSA – group level statistics for cluster: occipital cortex.* Grand-average RDM and maps of significant p-values are shown in Figure 2. I) testing vs. 0: FDR-significant results were obtained for 39 % (physR) of RDM fields, 72 % (joyR), and 80 % (physR vs. joyR). II) testing vs. shuffled (noise estimate): Permutation-significant results were obtained for 0 % (physR) of RDM fields, 22 % (joyR), and 44 % (physR vs. joyR).

### Supplement Figure 1) Rotation performance: Relative heading error.

The relative heading error (start-end; orientation yaw) is shown for all participants. The boxplot displays the median with whiskers extending to 1.5 times the interquartile range. Bonferroni-significant p-values of post hoc testing are shown (paired t-test). ** indicates p < 0.01. A) Relative heading error of the anti-clockwise rotation. B) Relative heading error of the clockwise rotation.
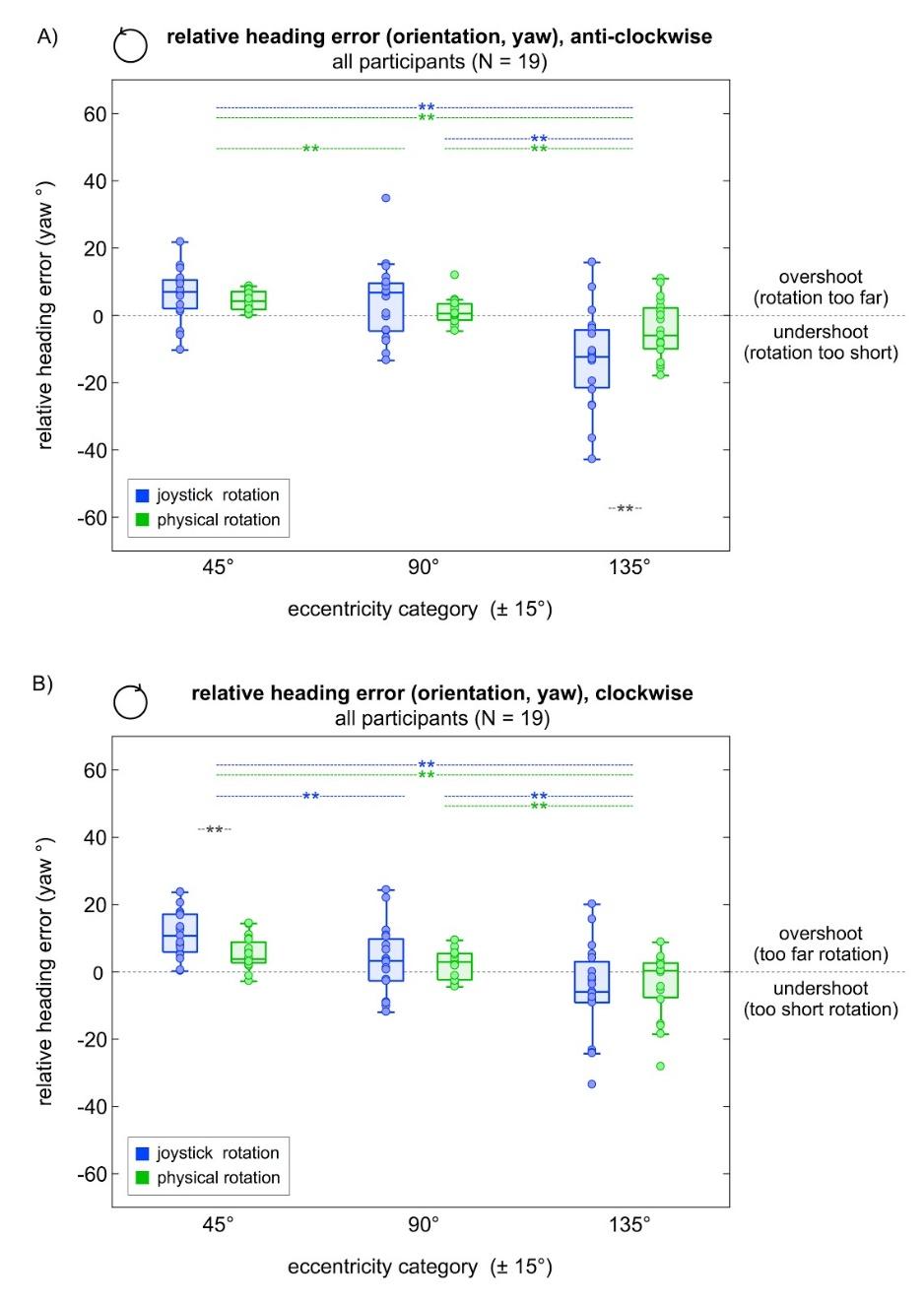

#### Supplement Figure 2A). Relative heading error, anti-clockwise rotation.

The relative heading error (start-end, orientation yaw) is shown for all participants. The boxplot displays the median with whiskers extending to 1.5 times the interquartile range.
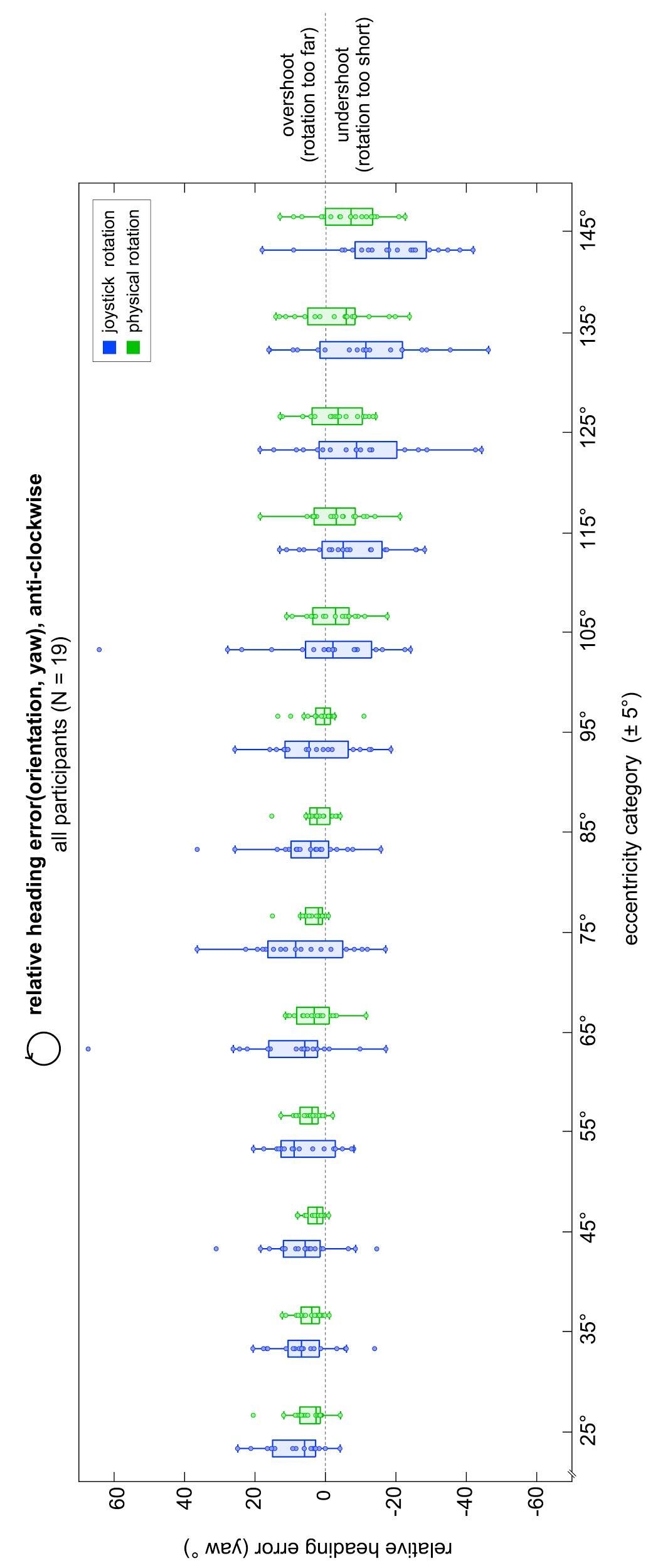

#### Supplement Figure 2B). Relative heading error, clockwise rotation.

The relative heading error (start-end, orientation yaw) is shown for all participants. The boxplot displays the median with whiskers extending to 1.5 times the interquartile range.
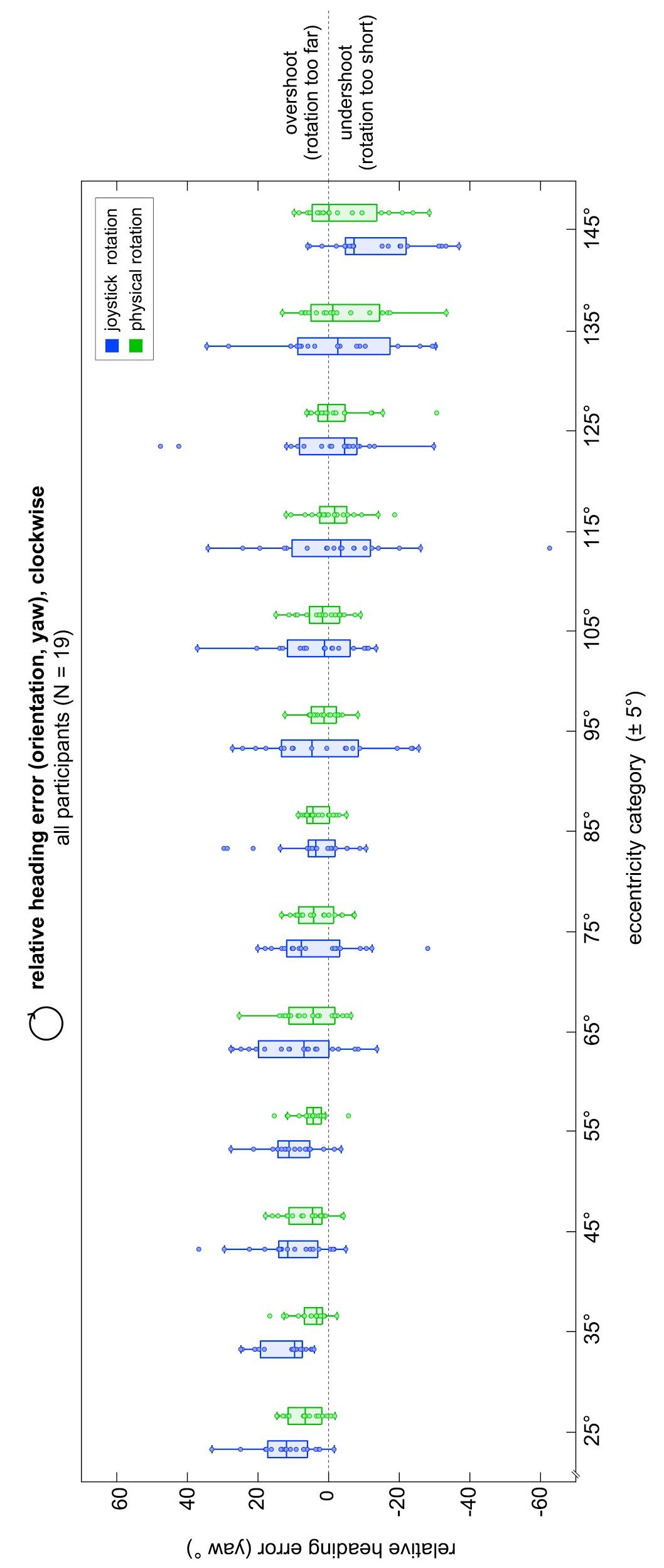

#### Supplement Figure 3) Reaction time for the outward rotation.

The reaction time is shown for all participants, indicated by the filled circles. The boxplot displays the median with whiskers extending to 1.5 times the interquartile range. ** indicates p < 0.01.
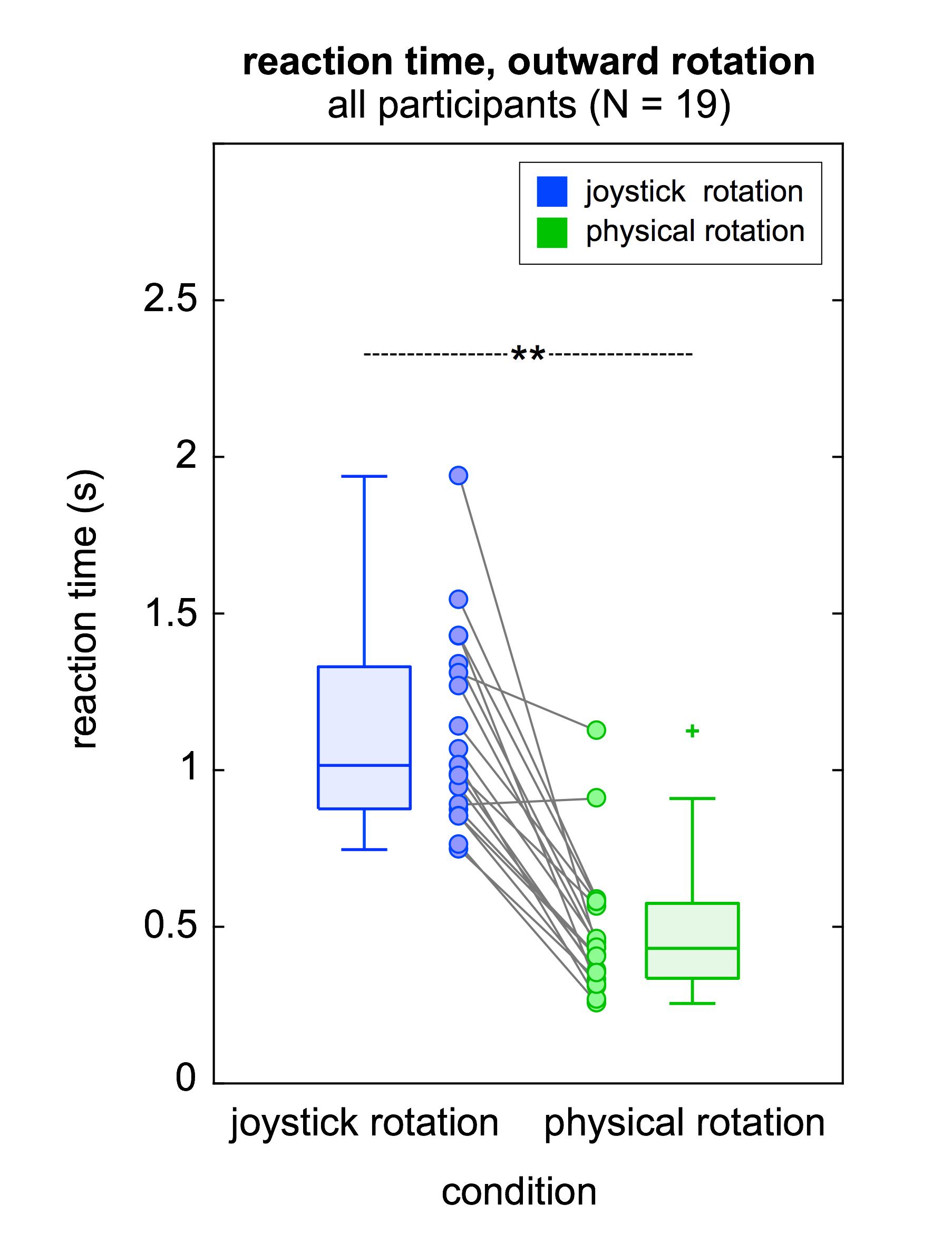

### Supplement Figure 4) Representational similarity analysis (RSA) of movement velocity-associated modulation of neural activity in single participants.

For five representative participants the IC topography (panels A-E) is shown (each IC belonging to the “right neck”, “eye”, “RSC”, “right parietal”, and “occipital” cluster, respectively). Panels F-J) show the baseline-corrected, trial-averaged amplitude of the respective IC activity per frequency band (ranging from 4-30.5 Hz) and movement velocity bin (10-100 % referring to slowest and largest velocities, respectively) during the outward rotation (time range between movement onset and offset). Displayed are the start categories of each frequency band (n=9 bands; non-overlapping 2.5 Hz steps). Each panel is scaled to its individual min. and max.; positive values indicate a relative increase of IC amplitude with respect to the pre-stimulus baseline, and vice versa for negative values. Panels K-O) display the RDM (180 x 180 entries) of the observed IC activity (Panels F-J), showing the normalized Mahalanobis distance (MD_norm_) for each bin x band combination. MD_norm_ = 0 indicates no distance and MD_norm_ = 1 indicates max. distance. Panels P-T) display the averaged RDM calculated on 1000 times shuffled IC activity (Panels F-J), with MD_norm_ ≈ 0.5. For visual inspection, the axes were clipped at 0.4 and 0.6.

IC – independent component; RDM – representational dissimilarity matrix; RSC – retrosplenial complex.

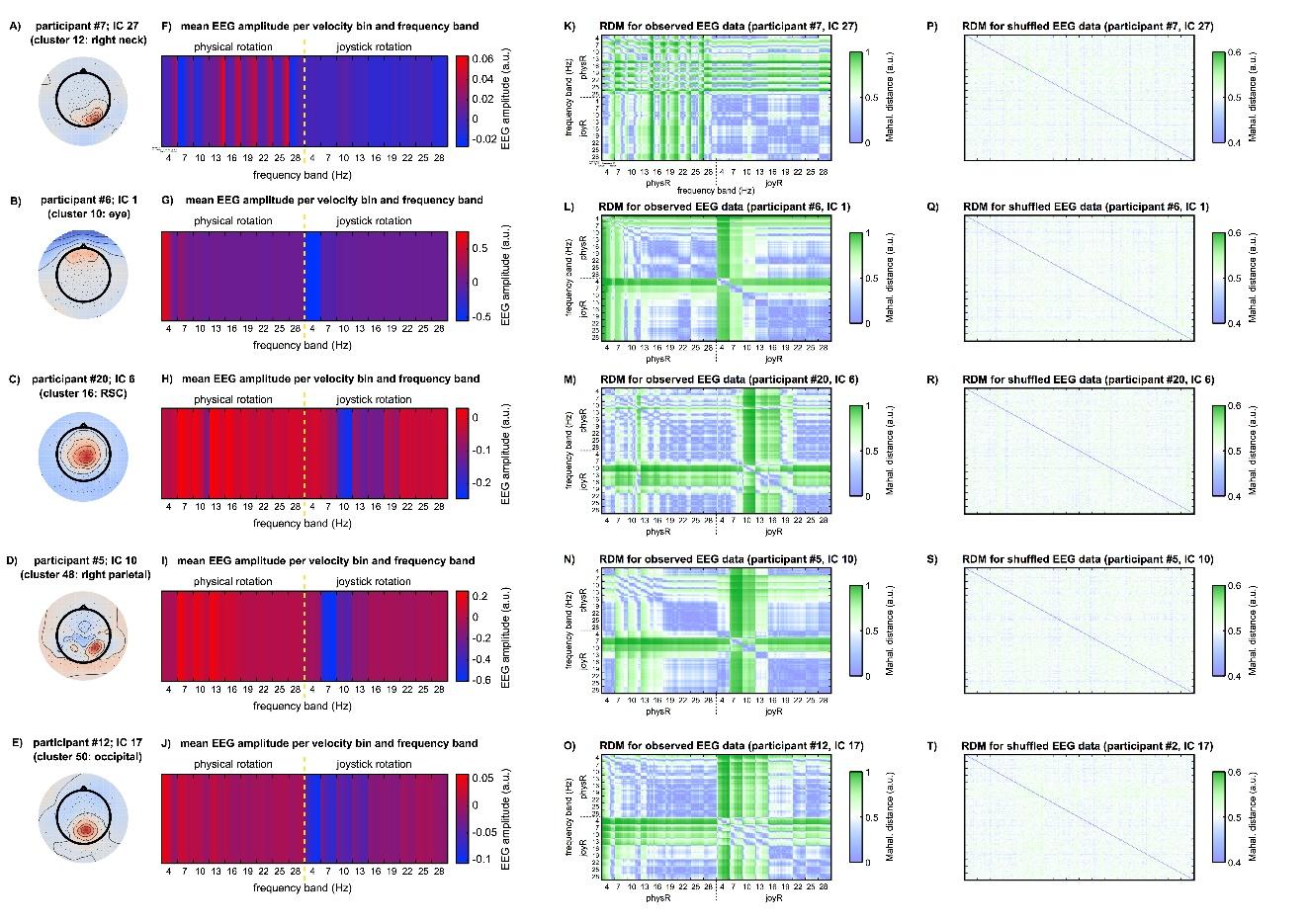
